# Detection of Pre-Existing Immunity to Bacterial Cas9 Proteins in People with Cystic Fibrosis

**DOI:** 10.1101/2025.03.20.644396

**Authors:** Gregory Serpa, Qiaoke Gong, Mithu De, Pranav S.J.B. Rana, Christopher P. Montgomery, Daniel J. Wozniak, Matthew E. Long, Emily A. Hemann

## Abstract

Cystic fibrosis (CF) is caused by homozygous mutations in the cystic fibrosis transmembrane conductance regulator (*CFTR*) gene, resulting in multi-organ dysfunction and decreased lifespan and quality of life. A durable cure for CF will likely require a gene therapy approach to correct CFTR. Rapid advancements in genome editing technologies such as CRISPR/Cas9 have already resulted in successful FDA approval for cell-based gene editing therapies, providing new therapeutic avenues for many rare diseases. However, immune responses to gene therapy delivery vectors and editing tools remain a challenge, especially for strategies targeting complex *in vivo* tissues such as the lung. Previous findings in non-CF healthy individuals reported pre-existing antibody and T cell dependent immune responses to recombinant Cas9 proteins, suggesting potential additional obstacles for incorporation of CRISPR/Cas9 technologies in gene therapies. To determine if pre-existing immunity to Cas9 from *S. aureus* or *S. pyogenes* was present or augmented in people with CF (PwCF), anti-Cas9 IgG levels and Cas9-specific T cell responses were determined from peripheral blood samples of PwCF and non-CF healthy controls. Overall, non-CF controls and PwCF displayed evidence of pre-existing antibody and T cell responses to both *S. aureus* and *S. pyogenes* Cas9, although there were no significant differences between the two populations. However, we observed global changes in activation of Th1 and CD8 T cell responses as measured by IFNγ and TNF that warrant further investigation and mechanistic understanding as this finding has implications not only for CRISPR/Cas9 gene therapy for PwCF, but also for protection against infectious disease.

## Introduction

Cystic fibrosis (CF) is a genetic disease hallmarked by a defective cystic fibrosis transmembrane conductance regulator (CFTR) protein that results in severe multi-organ disease manifestations with particularly damaging effects for the lung. While new highly effective CFTR modulator therapies (HEMT) have resulted in rapid reductions of disease burdens for people with CF (PwCF), there remains a significant population of PwCF who cannot benefit from HEMT. Globally, there are additionally major barriers restricting access to CF standards of care, including HEMT. Although HEMT is rightfully considered a breakthrough achievement to treat the underlying cause of CF, it is not a permanent cure for CF. Therefore, development of approaches to correct the underlying *CFTR* genetic mutations and restore CFTR function is a long-term goal of CF research. The discovery of the clustered regularly interspaced short palindromic repeats (CRISPR)/Cas9 system has resulted in new molecular tools to conduct precision gene editing and has rapidly transformed approaches to gene therapy. A recent example for successful application of these technologies is the first-FDA approved therapy that utilizes CRISPR/Cas9 genome editing technology in CD34^+^ hematopoietic stem cells to treat transfusion-dependent B-thalassemia or sickle cell disease is the^1^. This represents a major milestone in genetic therapies and demonstrates the potential utility of CRISPR/Cas9 genome editing approaches.

Many of the initial CRISPR/Cas9 genome editing tools utilized Cas9 proteins derived from bacterial species such as *Staphylococcus aureus* or *Streptococcus pyogenes*^*2,3*^, and the FDA approved treatment utilizes a synthetic guide RNA and a *S. pyogenes* Cas9 endonuclease^4,5^. As many people have either been colonized or exposed to *S. aureus* and *S. pyogenes*, it could be predicted that humans may harbor some level of adaptive immune responses to Cas9 proteins that could specifically target and eliminate cells subjected to and expressing this component of a gene editing vector. Indeed, recent discoveries showed that heathy adults harbored both antibody and cellular immune responses against recombinant Cas9 proteins^6,7^. Further, given that many people with cystic fibrosis (PwCF) have recurrent and chronic infections that start early in life^8^, especially with *S. aureus*, we hypothesized that PwCF may have enhanced pre-existing immunity to Cas9 proteins compared to a non-CF population. If accurate, this implies that there could be an enhanced barrier to effective utilization of CRISPR/Cas9 therapies in PwCF. However, there remains a general lack of understanding of the extent to which humoral and cellular adaptive immune cell responses are dysregulated in PwCF, which could impact this hypothesis. To test this hypothesis, we used ELISA based methods to evaluate the levels of anti-Cas9 IgG and assessed T cell populations, cytotoxic molecule expression, and cytokine production following Cas9 peptide stimulations of peripheral blood mononuclear cells (PBMC).

Here, we present results from a single-center study that evaluated the presence of pre-existing adaptive immune responses to *S. aureus* or *S. pyogenes* Cas9 in healthy controls and PwCF. Results from this study demonstrate similar IgG responses between PwCF and healthy controls, which supports the presence of pre-existing immunity to Cas9 proteins. We identified similar CD4 and CD8 T cell production of IFNγ and TNF in PwCF and healthy controls when stimulated with peptide pools from *S. aureus* or *S. pyogenes* Cas9. Additionally, we observed similar frequencies of granzyme B^+^ and CD107a^+^ CD4 and CD8 T cells, suggesting similar cytotoxic capabilities between T cells in PwCF and controls. To our knowledge, these results are the first demonstration of pre-existing immunity to Cas9 proteins in PwCF. These findings should guide inclusion of methods to evaluate pre-existing and development of immune responses to Cas9 proteins in future human gene therapy approaches in PwCF that utilize CRISPR-based gene editing technologies. Intriguingly, while no difference in the ability of CD4 and CD8 T cells to produce IFNγ and TNF following Cas9 or positive control CEF [cytomegalovirus (CMV), Epstein-Barr virus (EBV), influenza virus] peptides was observed, there was a significant reduction in the frequency of IFNγ^+^/TNF^+^ CD4 and CD8 T cells in PwCF compared to controls when stimulated with PMA/Ionomycin. FoxP3^+^ CD4 T regulatory cells were also significantly reduced in PwCF, confirming previously reported findings in the context of bacterial infection in PwCF^9,10^. Together, these results demonstrate that while there are no alterations in the Cas9 specific response in PwCF, there are broad defects in the ability of their CD4 and CD8 T cells to produce CD8/Th1 effector cytokines that warrant further study.

## Materials and Methods

### Human blood and PBMC Cryopreservation

Anti-coagulated peripheral blood was collected by venous puncture and plasma samples collected following centrifugation at 16,000 x *g* for 10 minutes at 4°C in Microtainer® Blood Collection Tubes (BD catalog # 365967). Blood from non-CF and people with CF was provided as de-identified biospecimens from the Cure CF Columbus Research Development Program Translational and Data Core under the NCH IRB and OSU IRB 2020H0399. Several additional non-CF samples were provided from OSU IRB 2014H0154. Plasma samples were stored at - 80°C until use for ELISA endpoint measurements. PBMC were isolated by gradient centrifugation utilizing Ficoll-Paque Plus (Sigma, catalog # GE17-1440-02) according to manufacturer’s instructions. 1 × 10^7^ total cells were resuspended in 90% HI-FBS + 10% DMSO and cryopreserved for storage in liquid nitrogen.

### ELISA reagents and protocol

Samples were processed in batches, where n=11 samples were thawed and used at the same time for assessment in four endpoint ELISAs for human serum albumin (HSA), tetanus toxoid, *S. aureus* Cas9, or *S. pyogenes* Cas9. ELISA methods were adapted from those previously reported^6^. *S. aureus* Cas9 (Sa Cas9) SaCas9 No NLS and *S. pyogenes* Cas9 (Sp Cas9) SpCas9 V3 without a nuclear localization signal (No NLS) were obtained from IDT.

Human Serum Albumin (HSA) was obtained from Sigma Aldrich (catalog # SRP6182) and inactivated tetanus toxoid was obtained from Cellero (catalog # 1002). Nunc Maxisorp ELISA plates (Fisher catalog # 12-565-135) were coated overnight at 4°C by dilution into 100 μL of carbonate-bicarbonate coating buffer (Sigma Aldrich, catalog # C3041-50CAP) to 0.5 μg of antigen per well for each of the recombinant Cas9 proteins, tetanus toxoid, or HSA. Following overnight coating, wells were washed 3 times with Tris buffered saline with Tween-20 (Sigma Aldrich, catalog # T9039-10PK), and plates were incubated with 200 μL blocking solution for 1 hour at room temperature. Blocker BSA (10%) from ThermoFisher (catalog # 37525) was diluted to 1% in PBS for the blocking solution. Sample dilutions were performed in blocking solution (1% BSA in PBS) and were plated in technical duplicates. Plates were washed three times following blocking and 100 μL of samples were added to each well and incubated at room temperature for 2 hours. Plates were washed three times, followed by incubation with 200 μL of diluted secondary antibody for 1 hour at room temperature. The goat polyclonal, goat anti-human IgG Fc secondary antibody (R&D Systems, catalog # NB7449) was diluted 1:100,000 in 1% BSA. Plates were washed an additional three times and 100 μL of room temperature development reagent was added to each well. Plates were incubated at room temperature protected from light; a timer was used to precisely track the development time of each plate and 50 μL of 1N sulfuric acid was added after 10 minutes to end the reactions. The absorbance at 450 nm was quantified using a Spectramax ID3 plate reader.

### PBMC Stimulation and Flow Cytometry

Peptide stimulations were performed as previously described^11^. Frozen PBMC were rapid thawed in complete RPMI containing 2μL benzonase (Sigma, catalog #706643), centrifuged at 350*g* for 10 minutes at 4°C, and resuspended in a 96-well U-bottom plate (5×10^5^ cells/well).

Media, defined *S. aureus*^*12,13*^ or *S. pyogenes*^*14*^ peptide pools (Table 1), or Peptivator CEF MHC Class I Plus positive control peptide pool (Miltenyi, catalog #130-098-426) was added to cells with brefeldin A (eBioscience, catalog #00-4506-51), recombinant human IL-2 (Sigma, catalog #11011456001) and anti-human CD107a AF700 (BD, catalog #561340) were added to wells and incubated at 37°C for 24 hours. Cell stimulation cocktail containing phorbol 12-myristate 13- acetate (PMA) and ionomycin (PMAI) (Invitrogen, catalog # 00-4970) was added to unstimulated wells for the final 6 hours of the incubation period. Following incubation, cells were labeled with Live/Dead Fixable Blue Dead Cell Stain Kit (Invitrogen, catalog #L34962) according to manufacturer instructions. Samples were then blocked with Human TrueStain FcX (BioLegend, catalog #422302) for 10 minutes at room temperature and subsequently surface stained with the following fluorescently conjugated antibodies for 30 minutes on ice: anti-human CD3 APC-H7 (BD, catalog #560176), anti-human CD4 BUV496 (BD, catalog #612936, anti-human CD8 BUV805 (BD, catalog #612889), anti-human CD45R0 BV570 (BioLegend, catalog #304226), anti-human CD45RA AF647 (BioLegend, catalog #305154). Samples were fixed and permeabilized with BioLegend True-Nuclear Transcription Factor Buffer Kit (catalog #424401) according to manufacturer’s instructions. Following permeabilization, samples were blocked as described above and stained with the following antibodies specific for intracellular antigens for 40 minutes on ice: anti-human FoxP3 Pacific Blue (BioLegend, catalog #320116), anti-human IFNγ BB700 (BD, catalog #566394), anti-human TNF BV750 (BD, catalog # 566359) anti-human granzyme B FITC (BioLegend, catalog #515403). Samples were washed and resuspended in PBS to run on a Cytek Aurora flow cytometer. Samples were analyzed using FlowJo (BD).

**Table 1.** Staphylococcus aureus and Streptococcus pyogenes peptides utilized for T cell stimulation.

|  | Peptide position | Peptide sequence | Citation |
| --- | --- | --- | --- |
| Sa Pool 1 | Sa Cas9_849-869 | DEKNPLYKYYEETGNYLTKYS | <sup>12</sup> |
|  | Sa Cas9_862-876 | GNYLTKYSKKDNGPV | 12 |
|  | Sa Cas9_918-935 | LDNGVYKFVTVKNLDVIK | 12 |
|  | Sa Cas9_936-951 | KENYYEVNSKCYEEAK | 12 |
|  | Sa Cas9_956-973 | ISNQAEFIASFYNNDLIK | 12 |
|  | Sa Cas9_964-972 | FIASFYNND | 13 |
| Sp Pool 1 | Sp Cas9_511-529 | VLPKHSLLYEYFTVYNELT | 14 |
|  | Sp Cas9_615-623 | ILEDIVLTL | 14 |
|  | Sp Cas9_615-629 | ILEDIVLTLTLFEDR | 14 |
| Sp Pool 2 | Sp Cas9_836-852 | RLSDYDVAAIVPQSFLK | 14 |
|  | Sp Cas9_883-898 | MKNYWRQLLNAKLITQ | 14 |
|  | Sp Cas9_988-997 | YLNAVVG TAL | 14 |
|  | Sp Cas9_1036-1051 | ATAKYFFYSNIMNFFK | 14 |

### Data analysis and statistics

Statistical analyses and calculations of the area under the curve were performed in GraphPad Prism Version 10.4.1. The specific statistical tests used are listed in individual figure legends. GraphPad Prism Version 10.4.1 was used for all statistical analysis.

## Results

### Blood IgG specific for Cas9 proteins is prevalent and equivalent in Healthy Controls and PwCF

To evaluate whether antibodies to recombinant Cas9 proteins exist in PwCF, an ELISA-based method was adapted for this study^15^. Samples were obtained from non-CF healthy individuals and PwCF (non-CF, median age 32.5 years, n=24, 79% female; CF, median age 24 years, n=35, 57% female). ELISAs were performed to detect relative levels of anti-*S. aureus* Cas9 (Sa Cas9) and anti-*S. pyogenes* Cas 9 (Sp Cas9) IgG utilizing anti-HSA IgG as a negative control, and anti-tetanus toxoid IgG as a positive control^6^. Based on the dilution curves, IgG levels between non-CF and CF are similar for each antibody specificity (Figure 1A). Calculation of the area under the curve (AUC) to quantify antigen-specific IgG levels (Figure 1B) revealed statistically significant increases in the levels of tetanus toxoid, Sa Cas9, and Sp Cas9 compared to HSA for non-CF controls and PwCF, although there were no significant differences when comparing non-CF controls or PwCF responses either Cas9 or the positive control. We performed an additional more stringent analysis by assessing if individual samples had positive antibody responses for Cas9 or tetanus toxoid defined by an AUC greater than 3 standard deviations above the mean AUC for HSA. As expected, antibody positivity to tetanus toxoid was high, with 89% of all tetanus toxoid samples having AUC values greater than then mean AUC of HSA plus three standard deviations, with little difference between non-CF healthy controls (87%) and PwCF (91%). The overall percent of samples antibody positive for Sa Cas9 was 57% and 68% for Sp Cas9. While minimal differences between non-CF healthy controls and PwCF for Sa Cas9, 58% versus 57%, respectively, were detected, 52% of samples from non-CF healthy controls were positive for Sp Cas9 while 75% of samples from PwCF were positive.

**Figure 1.**
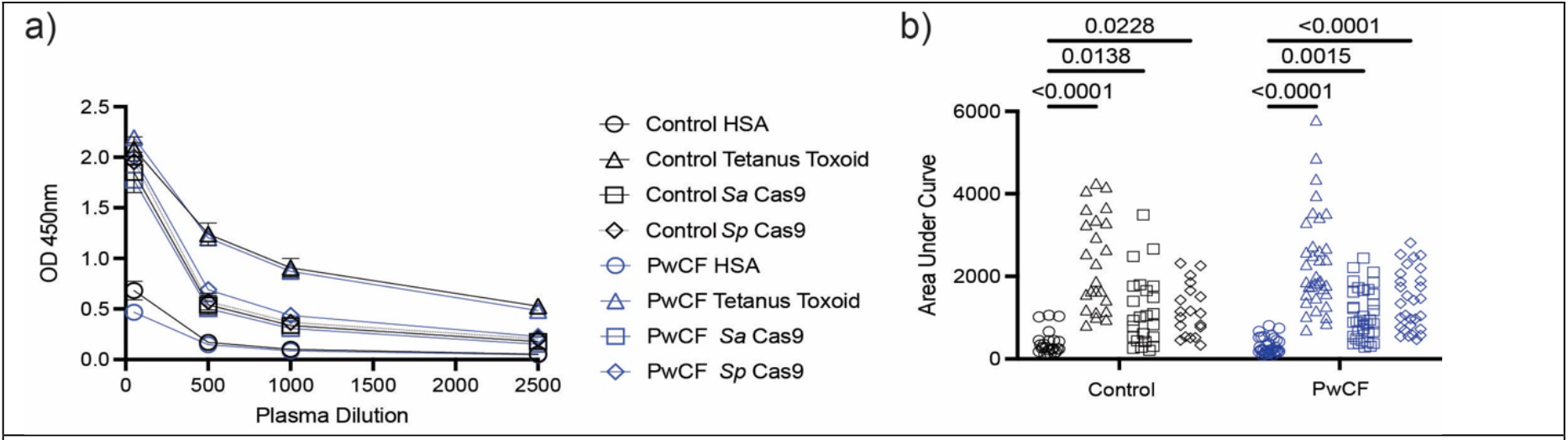
Detection of antibodies in blood of healthy donors and people with cystic fibrosis. (A) Mean ± SEM of ELISA measurements from one sample demonstrate OD450 nm endpoint values across sample dilutions of 1:50, 1:500, 1:1000, and 1:2500. (B) The area under the curve from the cumulative data obtained from all ELISA measurements for HSA (n= 28 non-CF control, n=34 PwCF), tetanus toxoid (n=28 non-CF control, n=34 PwCF), *S. aureus* Cas9 (Sa Cas9) (n=28 non-CF control, n=34 PwCF), *S. pyogenes* (Sp Cas9) (n=19 non-CF control, n=29 PwCF). Statistical comparisons were performed by two-way ANOVA with Tukey’s multiple comparison test; there were no statistically significant differences comparing non-CF control or PwCF for each individual ELISA.

While it is unclear what may account for this apparent increase in PwCF, it is perhaps due to a reduced sample size that was able to be performed for Sp Cas9. Overall, these results suggest that the prevalence of pre-existing immunity to Sa Cas9 and Sp Cas9 is relatively high in these two populations, consistent with previous findings reported in a cohort of healthy humans^6^.

### Cas9-specific cytotoxic molecules and IFNγ/TNF production by CD4 and CD8 cells are equivalent in Healthy Controls and PwCF

While antibodies are indicative of pre-existing immunity that may correlate to immune responses targeting Cas9 proteins or Cas9 gene-edited cells, cytotoxic CD8 and CD4 T cell responses could result in the functional targeting of gene-edited cells expressing Cas9 via MHCI and MCHII, respectively. Therefore, we next investigated T cell immunity in a small cohort of healthy controls (n=8, median age 39.5 years, 83% female) and PwCF (n=6, median age 31.5 years, 63% female). PBMC were stimulated with a number of previously identified *S. aureus* and *S. pyogenes* immunostimulatory Cas9 peptides, or CEF or PMA and ionomycin (PMA/I) as positive controls to determine if T cells from PwCF have altered responses specific for *S. aureus* or *S. pyogenes* Cas9 peptides compared to non-CF controls^12-14^. Following 24 hours of stimulation, flow cytometry analyses were performed to quantify the frequency of naive, memory, and regulatory T cells (Treg). Although T cell immunity in PwCF has not been widely reported in the literature, FoxP3^+^ CD4 Treg have been previously described to be significantly reduced in PwCF^16,17^. Specifically, reduced Treg responses in PwCF with chronic *P. aeruginosa* infection have been observed^10^. While the frequency of Treg was significantly increased *in vitro* following modulator treatment^9^, whether these levels compare to WT in PwCF on HEMTs is unknown. Despite the small cohort in our study, we confirmed that the frequency of peripheral Foxp3^+^ CD4 T cells was significantly reduced in PwCF compared to controls (Figure 2a). We also determined that the frequencies of CD45RA^+^ (generally naïve) and CD45RO^+^ (generally memory) CD4 (Figure 2b) and CD8 (Figure 2c) T cells in the peripheral blood revealed no dramatic changes between controls and PwCF, suggesting the distribution of naïve and memory T cell populations in the peripheral blood is unaltered in PwCF.

**Figure 2.**
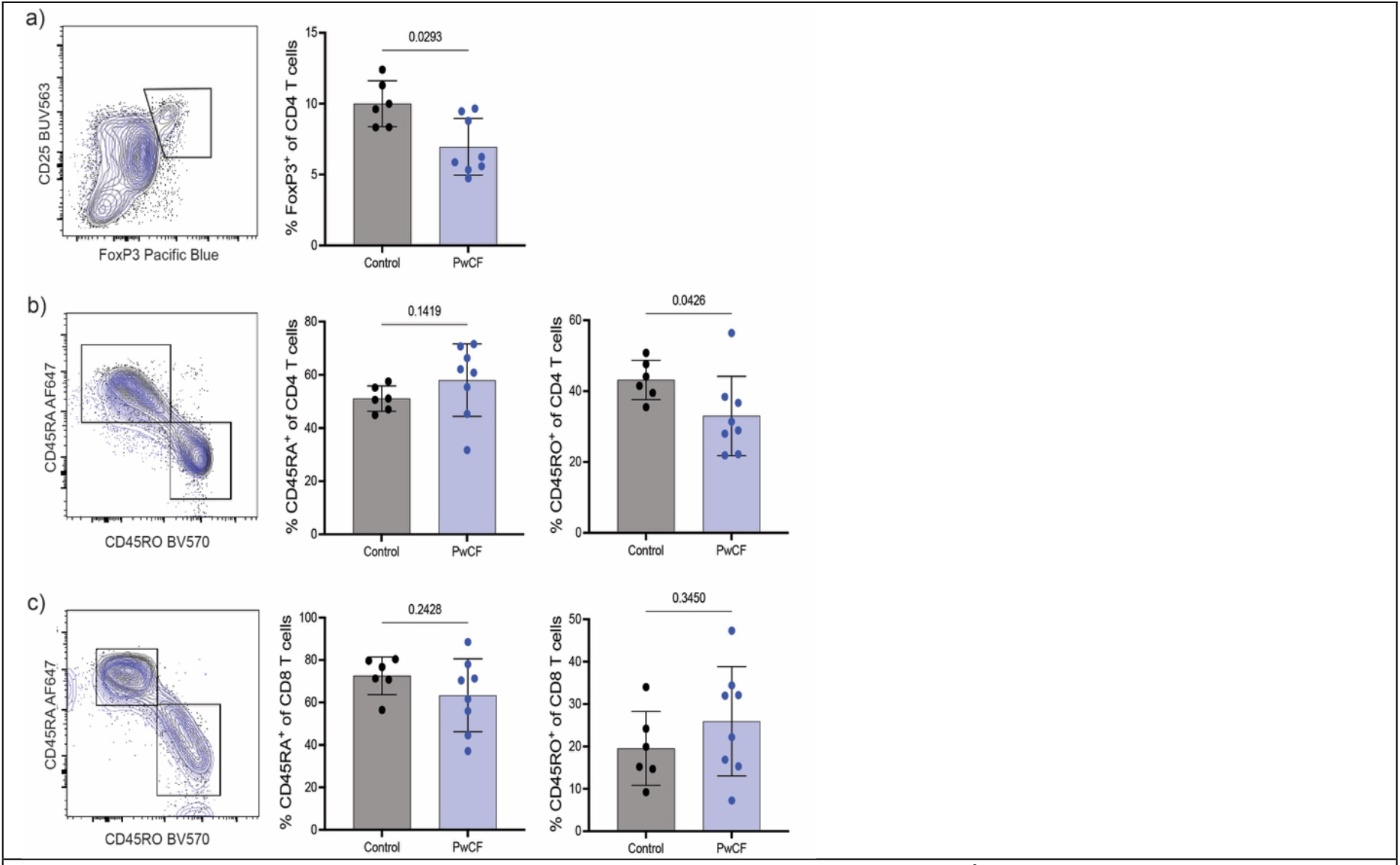
People with CF have decreased frequency of FoxP3^+^ CD4 Treg, but equivalent CD4 and CD8 T cell naïve and memory cell frequency compared to non-CF healthy controls. PBMCs were thawed and incubated in complete media containing IL-2 and brefeldin A for 24 hours, with PMA/I added for the last 6 hours. Cells were stained to identify (**a**) FoxP3^+^ CD4 T cells as well as CD45RA (naïve) and CD45RO (memory) CD4 (**b**) and CD8 (**c**) T cell populations by flow cytometry. Representative panels show gating of populations with healthy controls (black) and PwCF (blue) overlayed. Data points represent cells from an individual donor (n=6 non-CF controls and n=8 PwCF); bars represent the mean ± SD. Statistical comparisons were performed using an unpaired Student’s t-test.

To determine if the cytotoxic ability of CD4 and CD8 T cell responses was altered in PwCF, we measured the intracellular expression of granzyme B (gzmB) and found no significant differences in the percentage of gzmB^+^ CD4 (Figure 3a) or CD8 (Figure 3b) T cells. Additionally, we included fluorescently labeled anti-CD107a antibody during 6 hour stimulation with PMA/I in the presence of IL-2 and brefeldin A to serve as a proxy measure of cytotoxic degranulation as the antibody would bind CD107a expressed on the surface of cells during degranulation for analysis by flow cytometry^18^. The frequency of CD107a^+^ CD4 (Figure 3c) or CD8 (Figure 3d) T cells was not altered in PwCF, suggesting the degranulation ability of T cells is equivalent in controls and PwCF. Together, these findings demonstrate no alteration in cytotoxic molecule expression in PwCF compared to non-CF controls. As anticipated based on previous studies identifying the peptides utilized, low but detectable frequencies of CD4 T cells producing IFNγ and TNF were identified following stimulations (Figure 3e), although responses were largely undetectable following stimulation with the *S. aureus* (Sa) peptide pool. However, CD4 IFNγ/TNF production in response to both *S. pyogenes* (Sp) peptide pools was similar to the CEF positive control peptide pool, and responses were equivalent between control samples and PwCF (Figure 3e). Similar results were observed when we assessed IFNγ and TNF production by CD8 T cells following stimulation (Figure 3f). In sum, these results indicate that pre-existing effector CD8 and Th1 CD4 T cell responses specific to Cas9 can be detected and are equivalent between controls and PwCF. Critically, these finding suggest heightened pre-existing immunity to Cas9 in PwCF due to increased risk of bacterial infection and colonization may not be a barrier to development of gene therapy treatments for PwCF.

**Figure 3.**
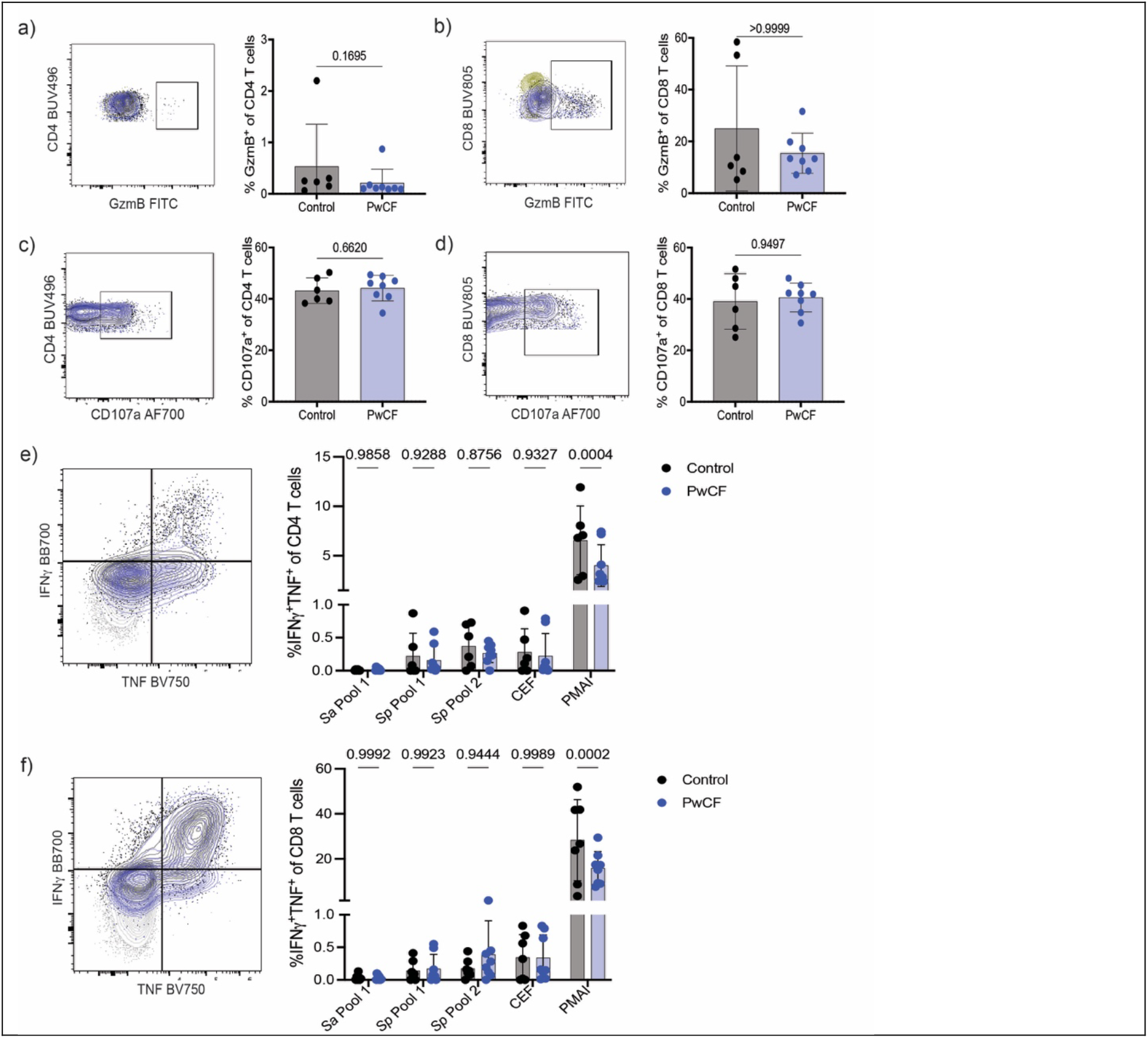
People with CF have equivalent responses to Cas9 peptides but a reduced total capacity for IFNγ/TNF production compared to non-CF controls. **a-d**) PBMCs were thawed and incubated in complete media containing IL-2, brefeldin A, and anti-CD107a for 24 hours, with PMA/I added for the last 6 hours. Cells were stained to identify the frequency of granzyme B^+^ CD4 (**a**) and CD8 (**b**) T cells as well as the frequency of CD107a^+^ CD4 (**c**) and CD8 (**c**) T cells. Representative flow plots show gating for populations of healthy controls (black) and PwCF (blue) compared to isotype control (yellow). **e-f**) PBMCs were thawed and incubated in complete media containing IL-2 and brefeldin A with Sp, Sa, or CEF peptides in Table 1 for 24 hours, or PMA/I added to the complete media for the last 6 hours. Cells were stained to identify the frequency of IFNγ^+^/TNF^+^ CD4 (**e**) and CD8 (**f**) T cells. Representative flow plots show gating for populations of healthy controls (black) and PwCF (blue) stimulated with PMA/I compared to baseline media incubation (grey). Data points represent cells from an individual donor from n=6 non-CF controls and n=8 PwCF and bars represent the mean ± SD. Statistical comparisons were performed using an unpaired Student’s t-test (**a-d**) or two-way ANOVA with Tukey’s multiple comparison test (**e&f**).

Despite observing no significant differences in the ability of CD4 or CD8 T cells from controls or PwCF to produce IFNγ/TNF in response to *S. pyogenes, S. aureus*, or CEF peptides, we noted a striking and significant reduction in the frequency of both CD4 (Figure 3e) and CD8 (Figure 3f) T cells producing IFNγ/TNF in PwCF following stimulation with PMA/I. This suggests that while the antigen-specific responses investigated are equivalent between controls and PwCF, the maximal effector response is specifically limited in PwCF. Whether these responses are blunted in response to infection or other stimuli still remains unknown. In summary, while this study indicates no additional risks for adaptive immune-mediated elimination of Cas9 targeted cells that would hamper gene targeting efforts in CF, we have identified a previously unreported defect in the ability of Th1 and CD8 T cells to produce IFNγ/TNF in response to PMA/I. These findings have potentially important implications regarding protective immune response to pathogens in PwCF that should be investigated more thoroughly in future studies.

## Discussion

CFTR modulator therapies have resulted in large advances in the quality of life for many people with CF. However, despite these advances, significant work remains to be accomplished to achieve a permanent cure for CF, regardless of *CFTR* mutation. The goal to provide a long-lasting cure for CF by correcting the underlying *CFTR* mutation has remained elusive since the discovery of the *CFTR* gene^19^. Immune responses to viral vectors used for delivery of gene therapies and limitations in the size of the cargo that can be packaged in different delivery system are challenges that have limited the ability to correct *CFTR* mutations^19^. However, new advancements in multiple technologies, including delivery vectors and genetic approaches of DNA replacement, mRNA replacement/repair, or gene editing, or premature termination codon (PCT) read-through agents/mRNA stabilizers are positioned to potentially provide new breakthrough therapies for PwCF^20^. Of all the approaches under development, gene editing is likely the best option, if perhaps the most challenging, to provide a durable cure for CF. Importantly, CRISPR/Cas9 gene editing technologies have resulted in rapid advancements in the ability to perform gene editing in cells *ex vivo* or *in vivo* ^21^. The CRISPR/Cas9 system was discovered in bacterial species, and different Cas9 proteins (the endonucleases that cut target sequences of DNA) that have been adapted as molecular biology tools are derived from bacterial species that include *Staphylococcus aureus* and *Streptococcus pyogenes* ^21^.

CRISPR/Cas9 gene editing technologies have ushered in new hope to provide gene therapies for a wide range of human diseases. Recent success utilizing *ex vivo* correction of Sickle cell disease is one example of an encouraging success. However, this clinical procedure requires a myeloablation regimen prior to delivery of edited cells and therefore has the potential advantage of reducing or eliminating immune responses to edited cells^22^. While the delivery of successful gene editing technologies has many hurdles, recognition by the host immune system is one component requiring further investigation as it is unknown whether pre-existing immunity to components of the CRISPR/Cas9 system already exists in PwCF. Based on surveys of non-CF populations ^6,7^, we predicted that pre-existing immunity in people with CF will be high.

Here, we performed the first assessment of pre-existing immunity to Cas9 proteins in PwCF. Results from this study revealed that consistent with non-CF individuals^6,7,12^, PwCF have correlates of pre-existing immunity to Cas9. These findings support the notion that that assessment of immune responses should be incorporated prior to and following delivery of any future CRISPR/Cas9 based genetic therapies. Additionally, identifying pre-existing immune responses in individuals may allow for development of modified Cas9 proteins to avoid these responses. In this regard, pre-existing immunity has already been the target of several studies that have attempted to silence these responses to improve gene targeting effectiveness^13,14^.

While limitations of this study include assessment of PwCF from a single clinical research center, we believe that these results may serve as a roadmap to incorporate these data into the efforts to find a permanent cure for CF.

In the current study, Cas9 peptide stimulations confirm the ability of PBMC from PwCF to respond even though no significant differences were observed compared to healthy control samples. Overall, while these results are consistent with the presence of T cell responses specific to Cas9 proteins reported in healthy controls^7^, one limitation of the current study is the small cohort size and lack of HLA phenotyping for the PMBC stimulations. This could contribute to some variability and potentially lower T cell (or PBMC) responses as we used peptide pools capable of stimulating MHCI and MHCII rather than screening to identify specific peptides associated with responses for specific HLA MHCI or MHCII types. While a larger cohort of individuals with defined HLA haplotypes would provide a more robust assessment of which *S. aureus* and *S. pyogenes* epitopes individuals are likely to be responsive to, our data demonstrate no significant changes in the frequency of CD4 and CD8 T cells expressing IFNγ and TNF. Additionally, the frequency of CD4 and CD8 T cells expressing CD107a and granzyme B are also equivalent between control PBMCs and those from PwCF. A further limitation of this experimental design is that the data only provide a surrogate of T cell function but did not directly assess the T cell mediated killing of target cells. Together, these results suggest there is no significant alteration in the cytotoxic T cell responses in PwCF that could potentially target cells that have been treated with CRISPR/Cas9 for CFTR mutation correction. However, the high prevalence of Cas9-specific responses suggest monitoring of these responses in conjunction with effectiveness of CRISPR/Cas9 gene correction therapeutics is warranted. Pre-existing immunity to CRISPR/Cas9 technologies may severely impact the success of any attempt to correct human *CFTR* mutations, and results from this study aim to help inform development of a long-lasting and durable cure for CF. Results from these experiments will have relevance to CF gene therapy approaches utilizing CRISPR/Cas9 technologies to correct CFTR mutations in cells in any organ afflicted by CFTR dysfunction.

While no significant alterations in Cas9-specific T cell responses were observed in this study, we did note global alteration in CD8 T cell cytokine production from PwCF. Following PMA/I stimulation, IFNγ/TNF producing CD4 and CD8 T cell frequencies were significantly reduced in PwCF. This suggests specific pathways downstream of PMA/I activation (including NFAT and NF-kB) may be defective and amplified by this stimulation compared to activation via antigen presentation by antigen presenting cells (APCs). Additionally, a recent study has identified airway fluid and neutrophils from PwCF to inhibit proliferation of T cell responses^23^. As our method of assessing cryopreserved PBMC samples would not allow for preservation of neutrophils, and we did not include airway fluid, it is possible that these factors may additionally impact Cas9-specific T cell responses in PwCF. Finally, while skewing towards Th2 CD4 T cell gene expression and limited Th1 gene expression has previously been shown to be associated with exacerbation and bacterial colonization^24^, the underlying mechanisms responsible for these phenotypes are not well understood and there remains a lack of specific investigation of potential alterations of the function of Th1 or CD8 T cell responses in PwCF. Future studies with larger cohorts will be critical to determine the full spectrum of defects in CD8 T cell responses in PwCF. Additional investigation of these adaptive immune responses will be critical not only for the development of gene therapies to correct CFTR function, but also to understand potential long-term consequences of altered immunity in PwCF with many living on HEMTs. Overall, additional studies investigating adaptive immunity in PwCF will be important to understand CF disease in PwCF in the context of both prolonged life expectancy due to HEMTs as well as the emergence of potential gene editing technologies.

## Acknowledgements

Recombinant Cas9 proteins were provided as a gift by Integrated DNA Technologies, Inc.

## Author Contributions

Gregory Serpa: Investigation, Writing-Review and Editing Qiaoke Gong: Investigation, Mithu De: Investigation, Pranav S.J.B. Rana: Resources, Writing-Review and Editing, Christopher P. Montgomery: Methodology, Writing-Review and Editing Daniel J. Wozniak: Resources, Writing-Review and Editing, Matthew E. Long: Conceptualization, Investigation, Visualization, Writing-Original Draft, Funding Acquisition, Emily A. Hemann: Conceptualization, Investigation, Visualization, Writing-Original Draft, Funding Acquisition. All authors have reviewed and approved the submitted document.

## Funding

Support for this work was provided by the Cystic Fibrosis Foundation awards LONG19F5-CI, LONG21R3 (M.E.L.) and Research Development (MCCOY19R0) Program Pilot and Feasibility Award (E.A.H.).

## Conflict of Interest Statement

There are no conflicts of interest to disclose.

## Data Availability Statement

Raw data is available upon request.

## Notes

### Competing Interest Statement

The authors have declared no competing interest.

